# Modelling FtsZ nucleation, hydrolysis and treadmilling activity with Monte Carlo methods

**DOI:** 10.1101/2020.02.25.965095

**Authors:** L. Corbin, H.P. Erickson

## Abstract

Bacterial cell division is tightly coupled to the dynamic behavior of FtsZ, a tubulin homolog. Recent experimental work *in vitro* and *in vivo* has attributed FtsZ’s assembly dynamics to treadmilling, where subunits add to the bottom and dissociate from the top of protofilaments. However, the molecular mechanisms producing treadmilling have yet to be characterized and quantified. We have developed a Monte Carlo model for FtsZ assembly that explains treadmilling and assembly nucleation by the same mechanisms. A key element of the model is a conformational change from R (relaxed), which is highly favored for monomers, to T (tense), which is favored for subunits in a protofilament. This model was created in MATLAB. Kinetic parameters were converted to probabilities of execution during single, small time steps, and these were used to stochastically determine FtsZ dynamics. Our model is able to accurately describe the results of several *in vitro* and *in vivo* studies for a variety of FtsZ flavors. With standard conditions, the model FtsZ polymerized and produced protofilaments that treadmilled at 28 nm/s, hydrolyzed GTP at 2.8 to 4.2 GTP min^-1^ FtsZ^-1^, and had an average length of 25 to 54 subunits, all similar to experimental results. Adding a bottom capper resulted in shorter protofilaments and higher GTPase, similar to the effect of the known the bottom capper protein MciZ. The model could match nucleation kinetics of several flavors of FtsZ using the same parameters as treadmilling and varying only the R to T transition of monomers.

**SIGNIFICANCE:** FtsZ assembly dynamics are now known to be governed by treadmilling, where subunits add to the bottom and dissociate from the top of protofilaments. We have generated a Monte Carlo model of treadmilling based on (a) a conformational transition of FtsZ subunits between two states, and (b) stochastic GTP hydrolysis. Importantly, the nucleation of new protofilaments is explained by the same mechanisms as treadmilling. We have determined kinetic parameters that match a wide range of experimental data. The model is available to users for their own *in silico* experiments.

## INTRODUCTION

Over the past two decades, research has cast FtsZ as a primary mediator of bacterial cell division. FtsZ is a homolog of eukaryotic tubulin, and like tubulin, it binds GTP and self-assembles long protofilaments (PFs) (1–4). Whereas tubulin PFs associate via lateral bonds to make the microtubule wall, FtsZ assembles single-stranded PFs under many *in vitro* conditions (4, 5). Early in the bacterial cell division cycle, PFs gather at the mid-cell forming a loose and highly dynamic ring, several PFs wide. This ring produces a constriction force on the membrane (6–8) and captains the voyage of the peptidoglycan synthesis machinery (9–11).

FtsZ exhibits cooperative assembly, as demonstrated by a sharp critical concentration (C_c_) and unfavorable nucleation (12–15). While multi-stranded filaments, such as actin, can achieve cooperative assembly by combining two types of bonds (16), the mechanism of cooperative assembly was not obvious for the single stranded FtsZ PF (5). Several groups subsequently suggested that cooperative assembly of a single-stranded polymer could be achieved if FtsZ had two conformations, one with high and one with low affinity for making the longitudinal bond (17–21). To explain cooperativity, the model proposes that the low affinity conformation is highly favored for FtsZ monomers, whereas the high affinity conformation is favored for subunits assembled in PFs. While some possible pathways encounter thermodynamic contradictions, Miraldi *et al*. (21) provided a detailed analysis of thermodynamically sound pathways. In short, the high affinity conformation alters both the top and bottom surfaces of the subunit to form a PF interface with a larger area of contact, and therefore with higher affinity, than the low affinity conformation. Even though the low to high affinity conformational switch is energetically expensive, if a subunit gains sufficient free energy from enhanced PF interfaces, the subunit will switch to high affinity when in a PF.

X-ray crystallography provided structural evidence for the two affinity states of FtsZ, which differ most obviously by rotation of the two globular subdomains. The N-terminal subdomain contains the GTP binding site and the top PF interface amino acids. The C terminal subdomain contains the bottom PF interface amino acids, including the conserved catalytic amino acids that trigger GTP hydrolysis when in a PF. Early crystal structures of FtsZ from multiple species, mostly monomeric, showed identical conformations, now interpreted as low affinity. In 2012, Matsui *et al*.and Elsen *et al*.crystallized *Staphylococcus aureus* FtsZ (*Sa*FtsZ) in the form of long, straight PFs (22, 23). The FtsZ in these PFs exhibited a striking conformational change relative to the previous monomeric forms. Noteably, the helix H7 moved downward one full turn, and the C-terminal subdomain rotated 25-28° relative to the N-terminal subdomain. More recently, two groups obtained crystals of *Sa*FtsZ with subdomain rotation corresponding to both the high and low affinity conformations (24, 25). Remarkably, both conformations assembled into PFs. However, the high affinity conformation showed a significantly larger interface surface area, 1,168 Å^2^, than the low affinity, 798 Å^2^(24). These two conformations have been designated T (Tense) for high affinity and R (Relaxed) for low affinity (24, 26). We will adopt this nomenclature.

Over the past 15 years, evidence has accumulated that FtsZ PFs treadmill, adding subunits at one end and losing them at the other. Early work demonstrated that FtsZ subunits exchange rapidly (half time approximately 8 s) between the Z ring and cytoplasm *in vivo* (27, 28) and between assembled PFs *in vitro* (29). Redick *et al*. found evidence that FtsZ PFs exhibit directionality by producing a collection of top and bottom interface capping mutants that inhibited assembly at either end (30). Mutations on the bottom interface blocked cell division while those on the top did not, suggesting that PFs assemble primarily at the bottom and disassembled at the top. Du *et al*. (31) confirmed this with mutants that exhibited a substoichiometric toxicity. The mutants in Redick *et al*. had an apparent weaker toxicity, but this may have been due to a fault in measuring the concentration of the mutant protein.

Recently, Loose and Michison (32) and Ramirez-Diaz *et al*. (33) observed and quantified treadmilling in swirling FtsZ vortices in vitro. Yang et al. (11) and Bisson-Filho et al. (10) reported similar dynamics in moving patches of FtsZ PFs in *Escherichia coli* and *Bacillus subtilis* cells. In each case, entire FtsZ patches moved across the membrane; however, single subunits within patches remained stationary, thus confirming treadmilling. Wagstaff *et al*. and Du *et al*. presented simple models of the treadmilling mechanism (25, 31).

The thermodynamics and kinetics of treadmilling were originally defined by Wegner (34). According to Wegner, treadmilling cannot occur for an equilibrium reaction. Treadmilling requires an irreversible step that alters the interface. For actin, this is ATP hydrolysis, and for FtsZ, this is GTP hydrolysis. Wegner also noted important thermodynamic constraints that must apply even to treadmilling generated by nucleotide hydrolysis.

In the present study, we designed a model for FtsZ treadmilling that combines the T and R state transitions with nucleotide hydrolysis, and satisfies Wegner’s constraints. With appropriate kinetic parameters, the model fits a wide range of experimental observations for FtsZ assembly dynamics.

## OVERVIEW OF DYNAMICS AND THE MODEL

FtsZ assembly dynamics can be divided into two main steps – nucleation and elongation. Nucleation is the creation of new PFs from free monomers. Once a PF is formed, monomers can freely associate and dissociate from the ends in a process called elongation. When elongation is directional, it produces treadmilling. In the following sections, we will first develop the model for treadmilling based on a combination of R and T state transitions, combined with GTP hydrolysis. We will then show how this same mechanism leads naturally to nucleation.

### Treadmilling mechanism based on R to T transitions and GTP hydrolysis

FtsZ monomers exist primarily in the relaxed R state, meaning that it requires a substantial positive free energy to transform a subunit to the T state. We assume that a subunit requires the same free energy to switch from R to T when it is incorporated in a PF. However, in a PF, a subunit gains a protein-protein interface. If the T conformation provides a larger and more stabilizing interface than R, this favorable negative free energy can compensate for the unfavorable R to T transformation.

Fig. 1 illustrates the R to T transition with geometric shapes representing the interfaces and conformational switch. The R subunits can assemble a PF, but the interfaces have gaps and the contact area is small, resulting in a weak longitudinal bond. If the subunits switch to T, the center arrow (meant to represent helix H7) moves down, opening its upper pocket to fit snugly with the full arrow above it. This movement also projects its own arrow point to create a new high affinity bottom interface. If the energy gained in the T-T interface, relative to R-R or T-R, is greater than the energy needed for the R to T transition of the subunit, subunits in a PF will favor switching to the T conformation.

**Figure 1:**
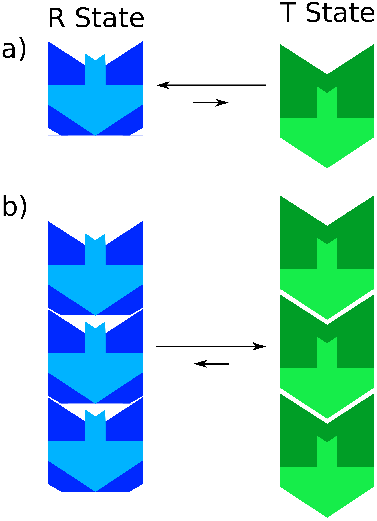
Illustration of the energetics of association related to the R and T conformation states. The R to T transition dominates assembly independent of GTP or GDP. We will later add parameters for GTP and GDP to modulate the association. **a)** The R-state is highly favored in monomers. **b)** When assembled in PFs the R-state produces interfaces with gaps. If the subunits transition to the T-state the interface becomes snug with a larger area. The larger interface area can compensate for the free energy needed for the R to T transition.

Fig. 2 shows the schematic model of how the two state system can generate treadmilling. We will describe the model by detailing the kinetics of each of the interfaces, labeled a-f in Fig. 2.

**Figure 2:**
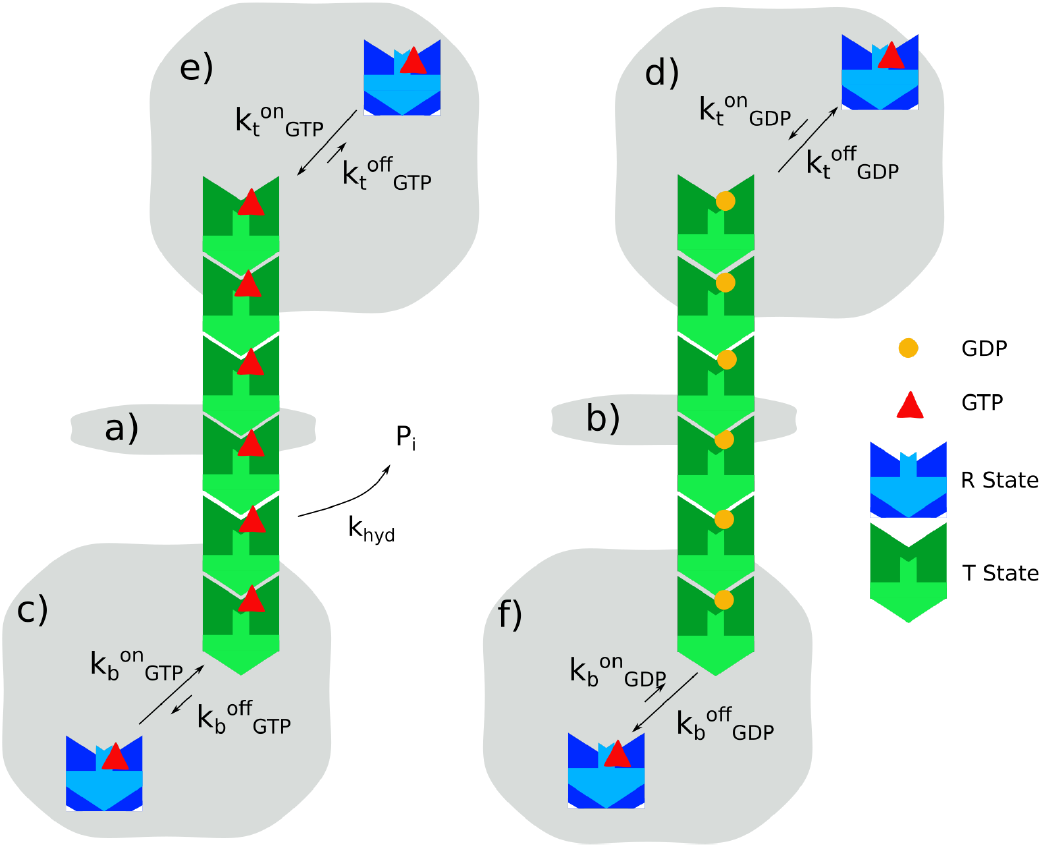
Our model for FtsZ treadmilling. Letters A-F refer to the interfaces whose kinetics are described in the text. The PF on the left has all subunits with GTP. The PF on the right has all GDP. The arrow in the center indicates the hydrolysis rate of a single GTP-bound subunit in a PF. Association is favored at the bottom of a PF when GTP is at the bottom interface. Dissociation is favored at the top of PFs when the top interface has a GDP.

a. This interface is in the middle of a PF and has a GTP. This interface is primarily stabilized by the subunits above and below the two subunits making the interface. The GTP further stabilizes it, making it the strongest interface. Our model assumes that the dissociation rate at this interface, which would result in fragmentation, is too slow to be observed during the time frame of the experiment.
b. This interface is similar to A, but has a GDP in the interface. We suggest that the GDP weakens the interface somewhat relative to interface A. But because the upper subunit is also sandwiched between two subunits, it is stabilized in the T conformation. This interface is still too strong to break during the time of the experiment, meaning fragmentation is negligible. Fragmentation and annealing could be added to the model later, but it has not been needed to explain treadmilling dynamics.
c. A new interface is formed when a monomer associates to the bottom of a PF with a GTP at the interface above it. This involves two steps, not shown explicitely. First, the subunit associates in the R state, which can occur with rapid, diffusion-limited kinetics. Then, the subunit has two pathways: it can dissociate, or it can convert to T. Because we have no data to distinguish these pathways, we combine them in a single step. k_b_^off^_GTP_ is the effective off rate assuming dissociation occurs before the transition to T. In the model, we can set the on and off rates arbitrarily, but once set, their ratio k_b_^off^_GTP_/k_b_^on^_GTP_ = K_D_ becomes a fixed parameter of the model. This will be essential for satisfying the Wegner constraints for interface E.
d. The top interface is the key to treadmilling. We postulate that when the top interface has a GDP, it is destabilized to the point that it switches to R and rapidly dissociates. The difference between this case and interface B (with GDP in a middle PF interface) is that the terminal subunit has no subunit above to stabilize its T-state. Like interface C, this reaction conceals two possible pathways. The primary path is switching to R and dissociating. Alternatively, the terminal subunit could exchange its GDP for GTP (since its GTP-binding pocket is exposed to solution) and then add another subunit. This would initiate a GTP cap at the top, discussed as interface E. We assume that the switch to R is much more favorable, but we include both possibilities by adding a hypothetical on rate. For thermodynamic equivalence, we include the possibility that the bottom subunit hydrolyzes its GTP, which lets it switch to R and dissociate similar to the top subunit. However, this is rare at the bottom relative to addition of a new subunit.
e. This interface is an essential pathway to accommodate the Wegner constraints. While there will be a gradient of high GTP at the bottom and high GDP at the top, the stochastic nature of hydrolysis means that there is a significant probability that a subunit will arrive at the top with a GTP in the interface below. In this case, the terminal subunit would be more stable than with GDP below. This terminal subunit with a GTP below can exchange its GDP for a GTP (since its GTP-binding pocket is exposed to solution), and then the addition of another subunit on top would be thermodynamically equivalent to adding a subunit to the bottom. This would initiate a GTP cap at the top and potentially compromise the treadmilling. However, the Wegner constraints offer an escape. They require that this addition be thermodynamically equivalent, which means that the dissociation equilibrium constant, K_D_, must be the same at the top and bottom. However, the kinetics can be slower at the top than at the bottom. Therefore, we can specify that k_t_^on^_GTP_ = k_b_^on^_GTP_/10 and k_t_^off^_GTP_ = k_b_^off^_GTP_/10. This keeps their ratio equal: K_D_ = k_b_^off^_GTP_/k_b_^on^_GTP_ = k_t_^off^_GTP_/k_t_^on^_GTP_. However, the slower kinetics at the top means that the new GTP cap on the top grows at 1/10 the rate at the bottom. Since GTP hydrolysis is constant, the top cap is eroded by hydrolysis faster than it elongates.
f. Interface F exists in the rare case that the bottom subunit in a PF has hydrolyzed its GTP. Because the interface does not contain a GTP to stabilize it and the bottom subunit is exposed, dissociation occurs rapidly. However, this case is rare because hydrolysis is significantly slower than addition of a new subunit.

### Intrinsic GTP hydrolysis and GTP turnover

GTP hydrolysis by FtsZ occurs only within a PF. Each subunit brings in a single GTP on its top when it associates on the bottom of a PF. Essential catalytic residues are provided by the T7 loop and helix H8 of the subunit above the interface (35–37). For our model, we assume that GTP hydrolysis occurs at a constant rate for any subunit within a PF. We call this the *intrinsic GTP hydrolysis rate*, and it is governed by the first order kinetic constant k_hyd_. Once a subunit has hydrolyzed its GTP, it retains the GDP until it dissociates from the PF and can exchange its GDP for GTP. Hydrolysis of a single subunit’s GTP is independent of hydrolysis in subunits elsewhere in the PF. PFs contain a mixture of GTP- and GDP-bound subunits. Since association occurs primarily at the bottom of a PF, this stochastic process produces a gradient of mostly GTP-bound subunits at the bottom and mostly GDP-bound subunits at the top.

Experimentally, GTP hydrolysis is measured as the release rate of either GDP or inorganic phosphate from the PF after GTP hydrolysis. This is different from the intrinsic hydrolysis rate, because GDP is released into solution only when a subunit arrives at the end of a PF and dissociates. We call this measurement *GTP turnover*. It is expressed as GTP min^-1^FtsZ^-1^. The GTP turnover is determined by a combination of k_hyd_, the concentration of polymer ends, and the kinetics of dissociation.

### Nucleation

We show next that the same processes that drive treadmilling provide a natural explanation for the nucleation of new PFs. Previously, nucleation was described as an unfavorable step to form a dimer, followed by much more favorable steps of elongation (15). The nature of the dimer was not specified, and it was actually difficult to reconcile with the single-stranded structure of the PF. We can now specify the pathway of nucleation in terms of the R to T transition, as shown in Fig. 3.

**Figure 3:**
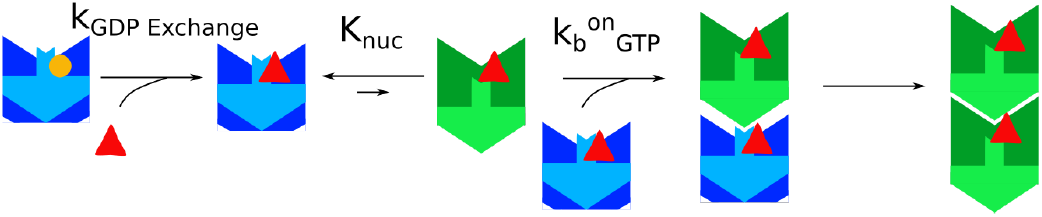
Our model for FtsZ PF nucleation. PFs nucleate from the rare free monomer that is in the T-state in solution. This T-state monomer can bind an R-state monomer with the same kinetics as a PF bottom. When that monomer switches to the T-state, a new PF is born. See Fig. 2 for key to symbols.

PF assembly is initiated experimentally by adding GTP to FtsZ monomers having bound GDP. The first step of nucleation must be activation of monomers by exchanging GDP for GTP, a first order reaction with rate constant k_GDP exchange_ (Fig. 3). In stop flow measurements, there was a 0.5 – 2 s lag that was independent of FtsZ concentration and was attributed to this nucleotide exchange (15). We assume an excess of GTP in the system, allowing for first order kinetics dominated by the dissociation of GDP.

The key step in our nucleation mechanism is the R to T transition of FtsZ monomers. The R state is highly favored for FtsZ monomers, but there must exist an equilibrium with a small fraction of T state monomers. We specify the equilibrium constant for this transition as [T]/[R] = 1/K_nuc_, where K_nuc_ is a large number indicating the unfavorability of the T-state in monomers. Our proposed nucleation mechanism suggests that a T-state monomer acts just like the bottom of a PF. R-state monomers can bind to it, and switch to T, with the same kinetics as they bind to the bottom of a PF. Once the T-T dimer is formed, it is considered a PF. A similar nucleation pathway was proposed by Dajkovic *et al*. (19); however, that model proposed that two monomers would switch to T and subsequently form the dimer nucleus. That pathway is equivalent in an equilibrium situation (21), but our proposal seems more reasonable kinetically. It has the additional attraction of using the same mechanism and kinetics as the elongation step in treadmilling.

Fujita *et al*. (24) suggested that the co-existence of R and T forms in the same crystal “indicated structural equilibrium of the two states either in aqueous solution or upon its crystallization.” Our model estimates the equilibrium ratio of monomers in solution, R/T, to be between 1,000 and 30,000, and as low as 10 for the mutant *Ec*FtsZ-L68W. This suggests that the structural equilibrium does not exist in solution, but occurs upon crystallization. We note that the PFs in the crystals can have subunits in either R or T state, but the area of the subunit interfaces are 798 and 1,168 Å^2^, respectively. If the crystal forms are isoenergetic, the extra interface energy of the T PFs should approximately equal the energy needed to transform an R subunit into T. We can check this with a simple calculation. A 10,000 R/T ratio corresponds to a free energy of 5.5 kcal/mol for the R to T transformation. The crystals were obtained at a protein concentration of ~ 100 μM FtsZ, which may approximate the K_D_ of the R interface, corresponding to ΔG = 5.5 kcal/mol. Adding 6 kcal/mol for the entropic free energy (16, 38), the ΔG_bond_ for the R PF interface would be 11.5 kcal/mol. Assuming that the interface bond energy is proportional to the interface area (39, 40), the T interface bond energy would be (1168/798) x 11.5 = 16.8 kcal/mol. The T interface is therefore 5.3 kcal/mol stronger than the R interface. This is very close to the 5.5 kcal/mol that we selected as the free energy needed for the R to T transformation. This is consistent with the suggestion that the extra interface energy of the T PFs is approximately equal to the energy needed to transform an R subunit into T.

**Figure 4:**
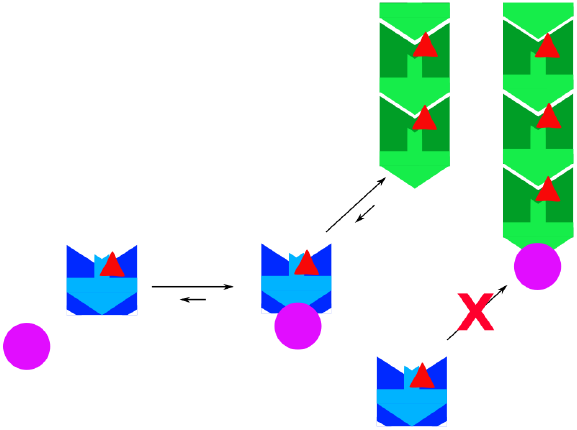
Capping mechanism of peptide MciZ (purple circle). MciZ binds the bottom of free FtsZ monomers in a reversible reaction. This complex can associate to the bottom of a PF. Additional subunits cannot associate below an MciZ molecule until the complex has dissociated. The FtsZ-MciZ complex dissociates rapidly if the FtsZ has hydrolyzed its GTP.

### FtsZ interacting molecules

A number of proteins interact with treadmilling PFs in the cell. Two classes that directly affect assembly are cappers and sequesterers. Cappers form a heterodimer with free FtsZ monomers, and these bind to the bottom of PFs and block future elongation. Sequesterers bind free monomers and prevent them from participating in the elongation and treadmilling reactions. In either case, our model assumes that the inhibitor rapidly establishes an equilibrium complex with FtsZ. A capper complex can bind the bottom of a PF with on-off kinetics that can be set by the user. For a sequesterer, we simply set to zero k_b_^on^_GTP_ and k_b_^on^_GDP_ for the capper-FtsZ complex.

### Lateral bonds

We want to emphasize that our model does not include lateral bonds. A recent study showed that ZapA causes increased PF bundling *in vitro* but did not alter treadmilling speed (41). Our model is consistent with this in assuming that treadmilling occurs at the level of single PFs and is not affected by their association into ribbons or bundles

## METHODS

We used Gillespie’s algorithm with partial equilibrium (42, 43) to model various steps of association and dissociation, GTP hydrolysis and nucleation. We developed the model in MATLAB R2019a (44), Mathworks Inc., Natick, MA. We used Matlab’s Parallel Computing Toolbox and the Duke Compute Cluster at Duke University for efficient data generation. Data analysis was performed in R, an open source collaborative project (45), with the packages matrixStats (46), dplyr (47), and beeswarm (48).

To run the model, the user initializes the simulation by setting kinetic parameters, the concentration of FtsZ and inhibitors, and the total time of the reaction. The model then calculates the integer number of molecules based on input concentrations for a volume of 2 fL, equivalent to that of an *E. coli* cell. We assume that the proportion of R to T monomers is at equilibrium and is not bound to an integer value, using partial equilibrium assumptions (43). The program then uses Gillespie’s algorithm to calculate the expected wait time until a single reaction occurs. The algorithm picks one reaction to implement based on the weighted probabilities. The current time is increased by the expected wait time. This process is repeated until it iterates over the entire time span.

To aid in the intuitiveness and accessibility of our model, we created a small app that can initialize and visualize the model. From this app, the user can also include bottom cappers or the mixing of two sets of PFs. The output has numerical and graphical displays of GTP turnover, monomer and polymer concentration as a function of time, as well as a graphic output showing the distribution of all PFs. The PF distribution can be replayed as a movie.

### Parameter optimization

In general, we estimated most kinetic parameters from experimental data; however, we did not have this sort of intuition for K_nuc_. To estimate K_nuc_, we performed parameter sweeps over a range of values and compared the initial assembly kinetics and Cc to experimental data.

To validate and further refine our choice of kinetics, we used simultaneous perturbation stochastic approximation (SPSA) (49). This algorithm searches for the best parameters to minimize the differences between the model output and experimental data. In short, the algorithm first implements small perturbations of the parameter sets in two directions and runs the model with both of the sets. For both parameter sets, the algorithm calculates the loss function, which is the difference between the model output and experimental data. These two points provide an estimate of the local gradient of the loss function. The algorithm then updates the parameters in the direction of the decreased loss. It repeats this process until it has reached a local minimum where the parameters fit experimental data the best. See jhuapl.edu/SPSA for resources on this method.

With SPSA, we optimized the kinetics to match experimental PF velocity, GTP turnover, and C_c_. We set the loss function to equal the mean absolute error of the model’s output compared to experimental data. Because the kinetics are many orders of magnitude apart, we simultaneously, randomly perturbed parameters by 1 to 10% in either direction and updated these each iteration. We iterated through this algorithm at least 50 times or until the loss function had reached a local minimum. If no decrease in loss was observed after 50 iterations, we experimented with starting parameters at a different point or by modulating the magnitude of the perturbations.

### PF nucleation and assembly

We measured assembly over time as the decrease in free monomers sampled each second for 15 to 30 seconds. Because our model is stochastic, it produces slightly different results each run. To portray our results, we show a shaded region giving the 95% confidence interval for the average of five runs. In these assembly curves, results from experimental assays are shown with dotted lines. (see Figs. 6d, 8a,b and 9a)

**Figure 5:**
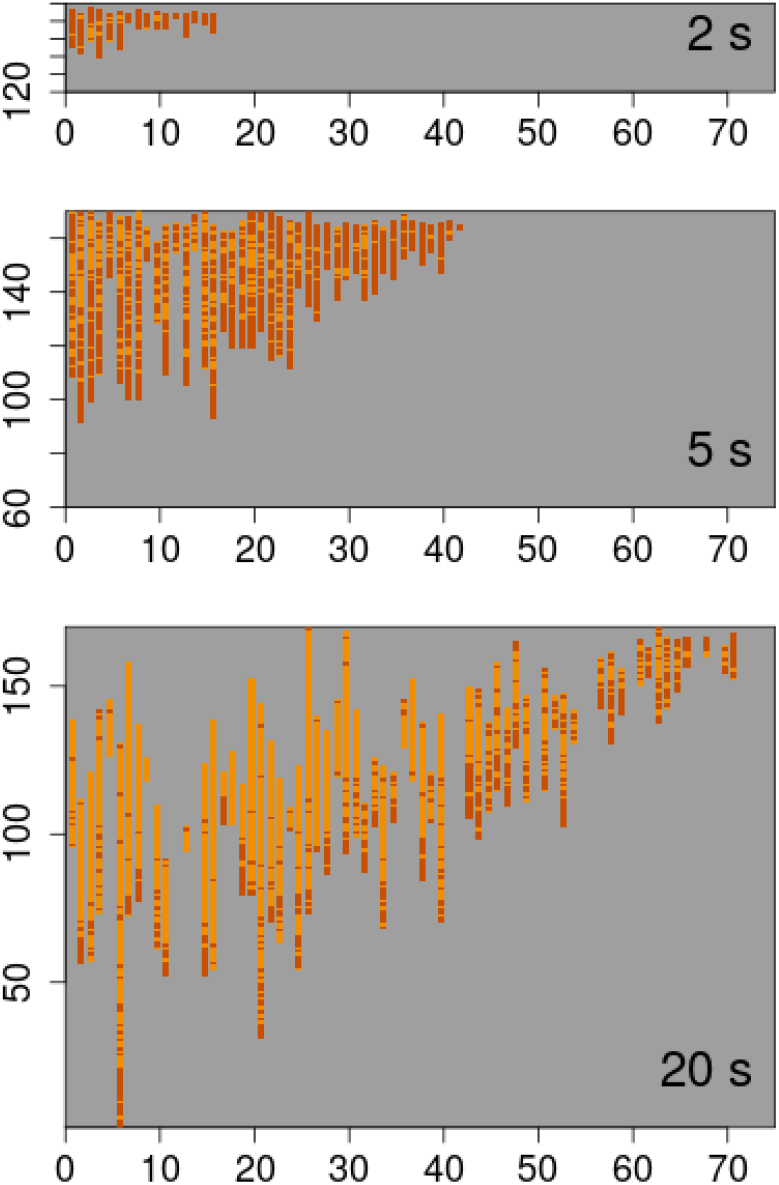
Visual representation of of the model’s output at 2, 5, and 20 s after initiating assembly. Each column represents a PF, shown as a stack of rectangular colored blocks, each representing an FtsZ subunit. The length of PFs in subunits is given on the y axis. Red represents GTP-bound FtsZ, and orange represents GDP-bound FtsZ. Newly nucleated subunits are added to columns at the top right.

**Figure 6:**
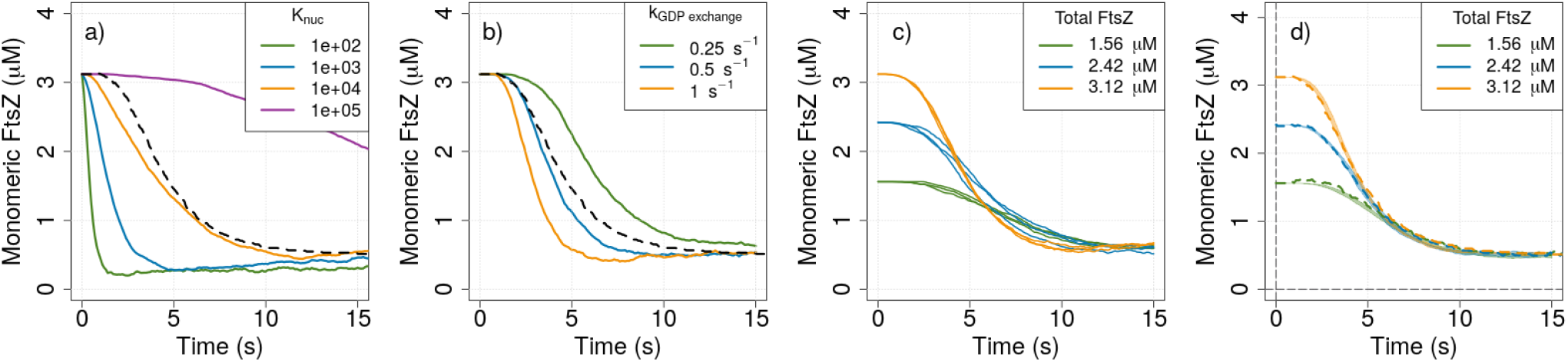
Nucleation of PFs from monomers over time. Assembly is shown as a decrease in monomeric FtsZ. We explored the two steps of nucleation separately and together with parameter sweeps attempting to match experimental kinetics of assembly. The black dotted line is experimental results for nucleation of 3.12 μM *Ec*FtsZ-F268C in MMK buffer, from Fig. 3 of Chen and Erickson. **a)** We first explored nucleation with only the rare R to T conformational switch over a range of K_nuc_. There was no nucleotide exchange step for these curves. **b)** Adding a step of nucleotide exchange improved the fit by introducing a lag.K_nuc_ was 1,000 for these curves. c) Demonstration of the stochastic nature of model. The model was run 3 times, each with three different concentrations of FtsZ with *Ec*FtsZ-F268C kinetics (tables 1, 2). **d)** Best fit of parameter sweep for nucleation of *Ec*FtsZ-F268C assembly. Shaded region is the 95% confidence interval of 5 runs, using the parameters for *Ec*FtsZ-F268C in Tables 1 and 2. Dotted lines are experimental results from Fig. 3 of Chen and Erickson.

To determine the C_c_, we measured the concentration of subunits assembled into PFs at steady state for a range of total subunit concentrations. To do this, we ran the assembly assays for one minute and measured the average PF subunit concentration over the last 20 seconds. (see Figs. 7a, 9b). The plot of assembled versus total FtsZ is a straight line, and C_c_ is the x-intercept.

**Figure 7:**
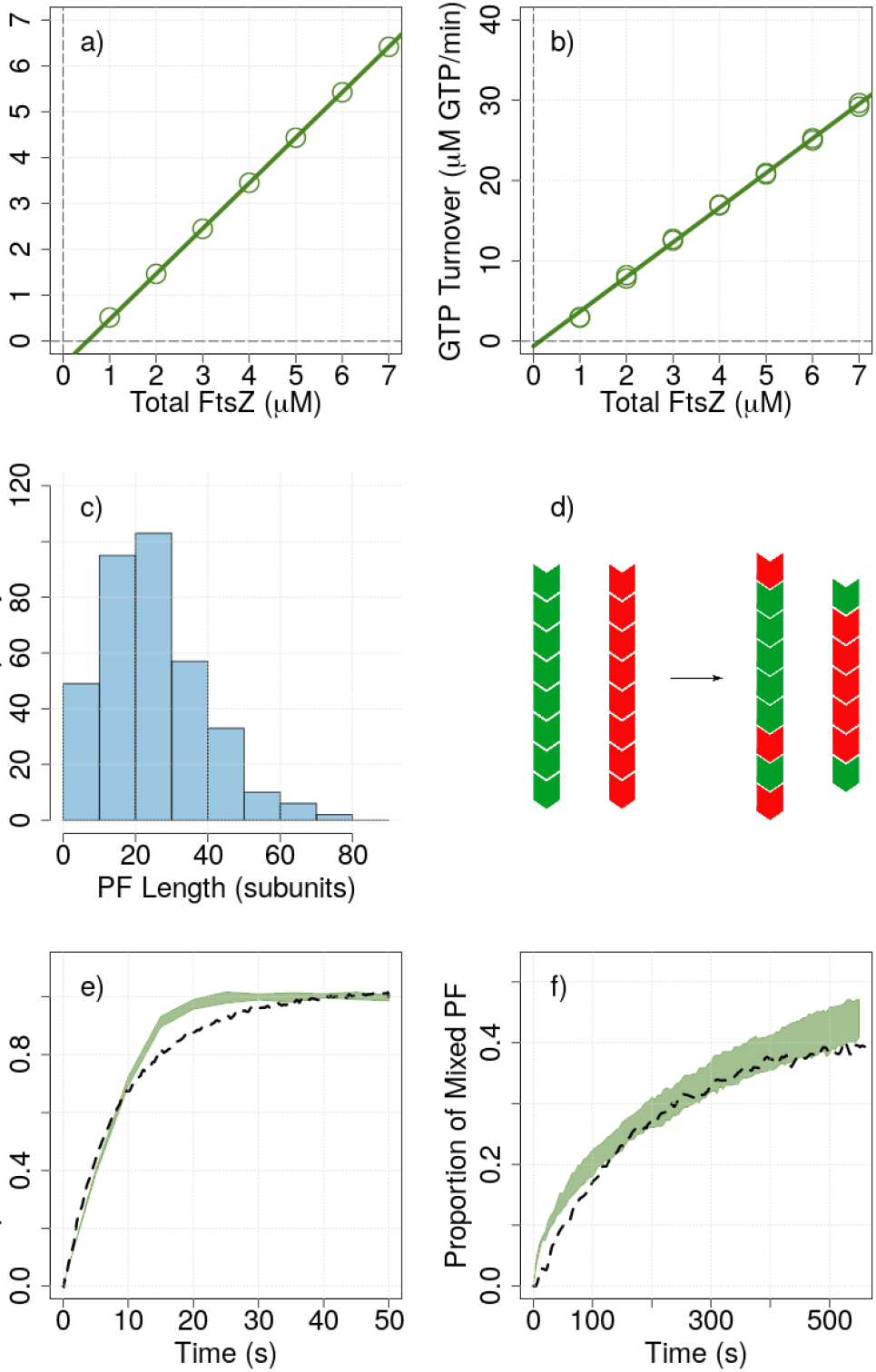
Summary of the model fit to experimental data for ***Ec*FtsZ F268C** (see Tables 1, 2 for parameters) **a)** Subunits in the PF vs. total FtsZ at steady state. The estimated C_c_ is 0.52 μM and slope is 0.99. **b)** Steady state GTP turnover as a function of total FtsZ. Avg. rate (slope) was 4.3 GTP min^-1^ FtsZ^-1^. The estimated C_c_ from GTP turnover was 0.14 μM. **c)** Steady state PF length with 2 μM FtsZ. (Avg: 25.0 ±13.9 subunits.) **d)** Diagram of the PF mixing experiment. Two sets of PFs are created – one with red dye and the other with green dye. These sets are mixed and the number of green-red interfaces and total interfaces are counted over time. **e,f)** Fit to inter-PF subunit exchange in the presence (e) or absence (f) of GTP hydrolysis. Zero indicates segregated PFs. One indicates mixed PFs. The dotted line is transformed experimental data from Chen and Erickson Fig. 4a (e) or Fig. 4b (f). Shaded region is 95% confidence interval of five model runs. The model was run with 6 μM FtsZ (3 μM each red and green).

### GTP turnover

GTP turnover has been measured experimentally by either the malachite green assay, which measures released phosphate, or a regeneration-coupled assay, which measures release of GDP. The latter assay is the one used for kinetics studies. The GDP is released only when a subunit dissociates, so we mimic this assay by counting GDP-bound subunits released by treadmilling. Experimental studies have generally measured the hydrolysis rate at steady state as a function of total FtsZ concentration.

To mimic these experiments, we ran our model for two minutes with a range of total FtsZ. We then determined the average release of GDP-bound subunits off the ends of the PFs over the last minute of the reaction. The x intercept of the linear regression of GTP turnover vs total FtsZ estimates a C_c_, independent of that determined from total polymer.

### PF length and velocity

At the end of every run, the program displays a histogram of PF lengths and calculates an average. The program also calculates average velocity of growth at the top, center, and bottom over the final five seconds.

### PF disassembly

To model disassembly of pre-assembled PFs, we used our model to first generate a set of PFs for a set amount of time. We then decreased k_b_^on^_GTP_, k_b_^on^_GDP_, k_t_^on^_GTP_, and k_t_^on^_GDP_ by 100 fold and continued the reaction until the PFs were fully disassembled. We measured the concentration of free monomer over time.

### Model availability

The Matlab code and documentation are available through the GitLab link: https://github.com/laurcor55/TreadmillModel. This includes a user friendly app that can access our model without coding knowledge.

## RESULTS AND DISCUSSION

### Default kinetic parameters

As indicated in Fig. 2, our model has 10 independent kinetic parameters and the equilibrium constant K_nuc_, which can be varied by the user. We will first discuss our rationale for default kinetic parameters (Table 1). Other kinetic parameters were fine tuned to fit specific mutants or species of FtsZ. These are given in the Table 2.

**Table 1:** Standard kinetic constants shared between all FtsZ and assembly buffers.

| Kinetic | Value |
| --- | --- |
| $k_b^{\text{on}}_{\text{GDP, GTP}}$ | $10 \mu\text{M}^{-1}\text{s}^{-1}$ |
| $k_b^{\text{off}}_{\text{GDP}}$ | $100 \text{s}^{-1}$ |
| $k_t^{\text{on}}_{\text{GTP}}$ | $1 \mu\text{M}^{-1}\text{s}^{-1}$ |
| $k_t^{\text{off}}_{\text{GDP}}$ | $7.5 \text{s}^{-1}$ |

**Table 2:** Kinetic constants unique to each mutant of FtsZ and assembly buffer.

| Protein | $K_{\text{nuc}}$ | $k_{\text{GDP exchange}}$ | $k_{\text{hyd}}$ | $k_b^{\text{off}}_{\text{GTP}}$ |
| --- | --- | --- | --- | --- |
| Ec F268C | 1,000 | $0.4 \text{s}^{-1}$ | $0.2 \text{s}^{-1}$ | $3 \text{s}^{-1}$ |
| Ec F268C (EDTA) | 10,000 | $0.4 \text{s}^{-1}$ | $0 \text{s}^{-1}$ | $3 \text{s}^{-1}$ |
| Ec L68W | 10 | $0.7 \text{s}^{-1}$ | $0.2 \text{s}^{-1}$ | $1 \text{s}^{-1}$ |
| Ec L68W (EDTA) | 1,000 | $2 \text{s}^{-1}$ | $0 \text{s}^{-1}$ | $3 \text{s}^{-1}$ |
| Bs | 30,000 | $1 \text{s}^{-1}$ | $0.5 \text{s}^{-1}$ | $8 \text{s}^{-1}$ |

**k_t_^off^_GDP_** = 7.5 s^-1^. This off rate is set to slightly higher than the experimentally observed treadmilling rate of 5-7 subunits/s in order to achieve rapid treadmilling. For completeness, we provide a **k_t_^on^_GDP_**. This must be rare in order to achieve directional treadmilling. We set k_t_^on^_GDP_ = 0.1 μM^-1^ s^-1^.

**k_b_^on^_GTP_** = **k_b_^on^_GDP_** = 10 μM^-1^s^-1^. At steady state, this balances the off rate at the top to create a C_c_ of 0.5-1.5 μM. 10 μM^-1^s^-1^ is near or above the generic diffusion-limited second order association constant (50). As explained above, we consider this a reversible reaction to account for the case where the incoming R subunit dissociates before it could switch to T. This rate helps drive the C_c_. We set rate constant, **k_b_^off^_GTP_**, between 1 and 8 s^-1^ so that the forward reaction is favored and to create the desired C_c_.

**k_b_^off^_GDP_** = 100 s^-1^. This is the off rate for the case where a bottom subunit hydrolyzes its GTP before an additional subunit associates below it. While it is rare to have a GDP-bound subunit on the bottom since GTP hydrolysis is slow, these kinetics were important for optimizing a rapid steady state GTP turnover.

**k_t_^on^_GTP_ = k_b_^on^_GTP_/10** and **k_t_^off^_GTP_ = k_b_^off^_GTP_/10** The left PF in figure 2 shows the case where the top subunit has a GTP below it. In this case, the GTP stabilizes the interface and the subunit remains in the T-state. It can initiate a GTP cap at the top if another subunit binds the top. This reaction has kinetics **k_t_^on^_GTP_** and **k_t_^off^_GTP_**. The GTP indicates a GTP in the top interface of the PF. As noted by Wegner (34), the equilibrium association constant for this reaction must be the same as that at the bottom, but we are free to vary the magnitude of the kinetics. We set kt^on^GTP = k_b_^on^_GTP_/10 and k_t_^off^_GTP_ = k_b_^off^_GTP_/10 to have a GTP cap at the top grow slowly and eventually be eroded by GTP hydrolysis.

**k_hyd_** = 0.2 to 0.5 s^-1^. This is the intrinsic GTP hydrolysis rate for the stochastic hydrolysis of any GTP with a subunit above it in the PF. k_hyd_ is a sensitive parameter and is the driving force behind many of our optimizations. To simulate an experiment with blocked GTP hydrolysis (assembly in EDTA), we set k_hyd_ to 0.

**k_GDP exchange_** = 0.4 to 2 s^-1^. This is the first order rate for a monomeric subunit with a bound GDP to exchange it for GTP, assuming excess GTP. Chen *et al*. and Chen and Erickson (15, 29) found that when assembly was initiated by adding GTP, it showed a lag of 0.5-2 s that was independent of FtsZ monomer concentration; they attributed this lag to the time for nucleotide exchange. We get a better fit to experimental nucleation by incorporating this parameter. We fine tune this parameter to fit specific nucleation rates.

Because the top of a PF is exposed to solution with a high GTP concentration, it would be able to exchange its GDP for GTP. However, during the dissociation from the top, we assume that the T to R transition is much faster than the time for nucleotide exchange. We chose not to include this reaction because it does not significantly influence the R to T transition at the top, as this is dependent on the penultimate subunit in our model.

**K_nuc_**= 10 to 30,000. This is the equilibrium constant for the R to T transition of monomers. This sensitive parameter was inspired by Miraldi *et al*. to produce cooperative assembly (21). K_nuc_ was optimized for each flavor of FtsZ by a parameter sweep. Species of FtsZ with a lower K_nuc_ have a lower C_c_ and are more isodesmic in nature.

The default kinetic parameters, listed in Table 1, were used as the starting point for all fittings. In most cases, the optimal fittings were achieved by varying K_nuc_, k_hyd_ and k_b_^off^_GTP_ with values listed in Table 2. The effect of varying other parameters are discussed below when needed.

### Initial exploration of nucleation of *Ec*FtsZ-F268C

Chen and Erickson (29) used the mutant *Ec*FtsZ-F268C, fluorescently labeled for FRET, to explore several aspects of assembly. We first show how our model was fit to the kinetics of initial assembly, which are determined by nucleation.

In our final model, there are two steps in nucleation: activation of the subunit by exchanging GDP for GTP, and the association of an R monomer to a (rare) T monomer to form a T-T dimer. We first explored nucleation using only the R to T switch and then integrated an activation step of GDP to GTP exchange.

The concentration of rare T monomers is defined by the equilibrium constant K_nuc_: [T]/[R] = 1/K_nuc_. A parameter sweep of K_nuc_ revealed a close match to experimental results when K_nuc_ was on the order of 1,000 (Fig. 6a); these curves were run without GDP.

We next added a GDP to GTP exchange step with the kinetic constant k_GDP exchange_ (Fig. 6b). Using the K_nuc_ = 1,000 determined in Fig. 6b, a k_GDP exchange_ of 0.4 s^-1^ improved the fit by introducing an initial lag in assembly.

To show the stochasticity of our model, we ran the model three times and showed the assembly of PFs over the first 15 seconds. The model gave slightly different results each time it was ran (Fig. 6c). For subsequent results, we ran the model five times for each condition and presented a shaded curve representing the 95% confidence interval of the average (Fig. 6d). Fig. 6d shows our fit to the experimental data of Chen and Erickson for nucleation of *Ec*FtsZ-L268C in MMK buffer, using the optimized parameters. The parameters for this fit are given in Tables 1 and 2.

### Fitting C_c_ and PF mixing of *Ec*FtsZ-F268C

We have shown the fit to nucleation kinetics in Fig. 6, as discussed above. We next used the model to fit the C_c_, PF length and a mixing experiment.

*Ec*FtsZ-F268C has a C_c_ of 0.5 μM, which is close to wild type (experimentally determined in MMK pH 6.5 buffer). Experimentally, there are two measures of C_c_. The first plots total subunits in polymer against total FtsZ. As shown in Fig. 7a, this gave a straight line with x-intercept at 0.52 μM FtsZ, close to the experimental C_c_ of 0.5 μM (29).

The second estimate of C_c_ is to plot GTP turnover as a function of total FtsZ (Fig. 7b). This also gave a straight line with a slope of 4.3 GTP min^-1^ FtsZ^-1^. This is close to the experimental value of 4.5 GTP min^-1^ FtsZ^-1^. The x-intercept of this plot gave a C_c_ of 0.14 μM, which is less than the C_c_ measured by assembly (Fig. 7a,b). We remain uncertain for this discrepancy.

The model gave a distribution of PF lengths 20 s after assembly shown as a histogram in Fig. 7c. The predicted average of 25.0 ±13.9 subunits is within the EM measurements of 20 to 40 subunits (29).

Finally, we used our model to fit the FRET data for exchange of subunits between PFs (29). Subunit exchange was measured experimentally by assembling separate pools of PFs labelled with either green or red dye. The PFs were mixed, and exchange was measured by the decrease in donor FRET signal (29). To model this, we preassembled two pools of FtsZ and labeled the subunits G for green and R for red, each with uniformly labeled PFs treadmilling in a pool of monomers at concentration C_c_. We then mixed the two pools and tracked the mixing. FRET, measured experimentally by donor quenching, occurs when a G and R subunit are adjacent in a PF, but a G with R on each side counts as only one. For example, GGRGRR counts as two G’s mixed. Our model gave a reasonably close match to the experimental data (Fig. 7e). Mixing in the absence of GTP hydrolysis was much slower but was also fit by the model after setting k_hyd_ to zero and K_nuc_ = 10,000 (Fig. 7f).

### Fitting nucleation, PF length and disassembly of *Ec*FtsZ-L68W

We next examined the dynamics of *Ec*FtsZ-L68W. This mutant has a significantly lower C_c_ than *Ec*FtsZ-F268C. When we performed parameter sweeps, we found K_nuc_ of 10 gave the best fit to experimental data (fig. 8a). While this K_nuc_ is low compared to other flavors of FtsZ that we examined, it agrees with the fitting of *Chen et al*. where the K_D_ for the dimer was only 12-fold higher than that for elongation (15). This confirms that the L68W mutation enhances the subunit interface and assembles with kinetics far from wild type cooperative assembly.

**Figure 8:**
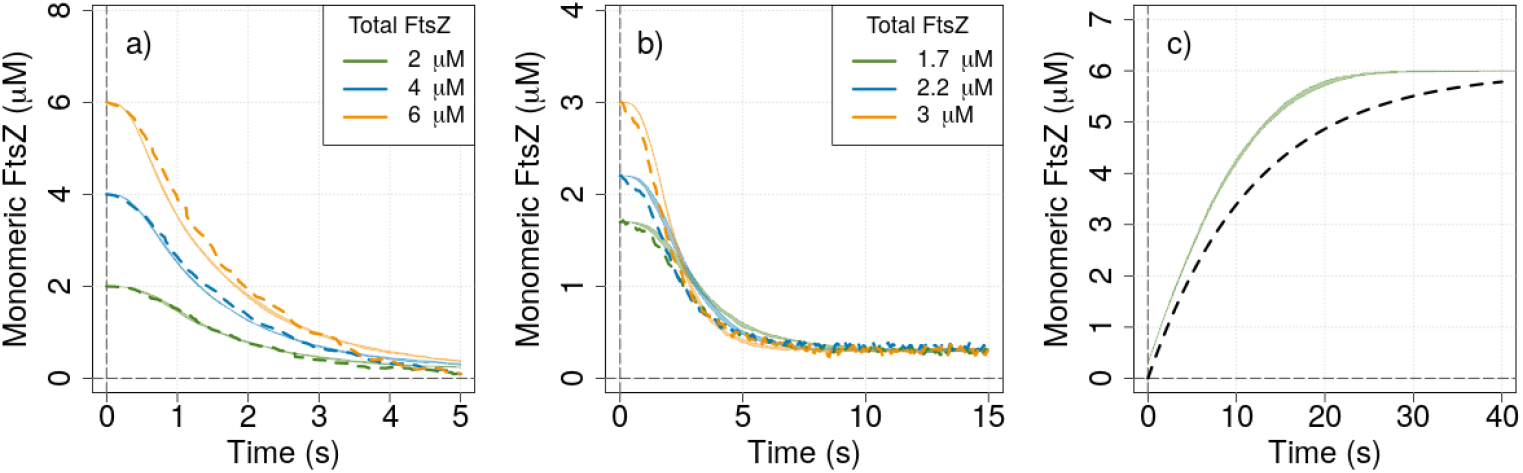
Summary of results with ***Ec*FtsZ-L68W**. **a)** Assembly of monomers over time in HMK buffer (k_hyd_ is set to 0.3 s^-1^). The model was initialized with standard kinetics and K_nuc_ = 10. Shaded region is the 95% confidence interval of 5 runs. Dotted lines are experimental results from Chen *et al*. Fig. 6c. **b)** Assembly of monomers over time in MEK buffer, where EDTA chelates Mg and blocks GTP hydrolysis (k_hyd_ is set to 0 s^-1^). Model was initialized with standard kinetics and K_nuc_ = 1,000. Dotted lines are experimental results from Chen *et al*. Fig. 6b. **c)** Disassembly of PFs after 20 s assembly period with 6 μM FtsZ. Shaded region is the 95% confidence interval of 5 runs. Dotted line is experimental data from Chen and Erickson Fig. 7c.

We next fit the nucleation of *Ec*FtsZ-L68W in MEK pH 6.5 buffer, where EDTA blocks GTP hydrolysis, by setting k_hyd_ to zero. The best fit is shown in Fig. 8b. The major adjustment was to increase K_nuc_ to 1,000. This is consistent with the original fit of Chen *et al*. where the K_D_ of the dimer was 400-fold higher than that for elongation (15). A similar adjustment of K_nuc_ was also used to fit the subunit exchange of *Ec*FtsZ-F268C in EDTA buffer (Fig. 7f).

To further examine *Ec*FtsZ-L68W dynamics over time, we performed a disassembly experiment that mimicked that of Chen and Erickson Fig. 7c (51). We used our model to pre-assemble PFs for 20 seconds. We then decreased k_b_^on^_GTP_ and k_t_^on^_GTP_ by 100 fold and allowed the reaction to continue until the PFs completely disassembled. We found this produced PFs that disassembled with a half-time of 5.7 seconds, as compared to 8.3 seconds found experimentally (Fig. 8c). This fit used only the parameters previously used to fit initial assembly, with no adjustment to better match the disassembly.

### Assembly of *Bs*FtsZ and effects of bottom capper MciZ

We then attempted to fit the extensive experimental data on FtsZ from *Bacillus subtilis, Bs*FtsZ. *Bs*FtsZ has a C_c_ between 1 and 1.5 μM (52). To achieve this C_c_ and match the nucleation kinetics, we performed a parameter sweep of K_nuc_ and k_hyd_. We found the best fit to be with K_nuc_=30,000 and when k_hyd_=0.5 s^-1^ (Fig. 9a). Furthermore, we increased the k_b_^off^_GTP_ to 8 s^-1^ to obtain the correct C_c_ (Tables 1, 2). The experimental data showed an assembly overshoot at 10-15 s. Our model did not match this overshoot but did match the plateau at >25 s.

**Figure 9:**
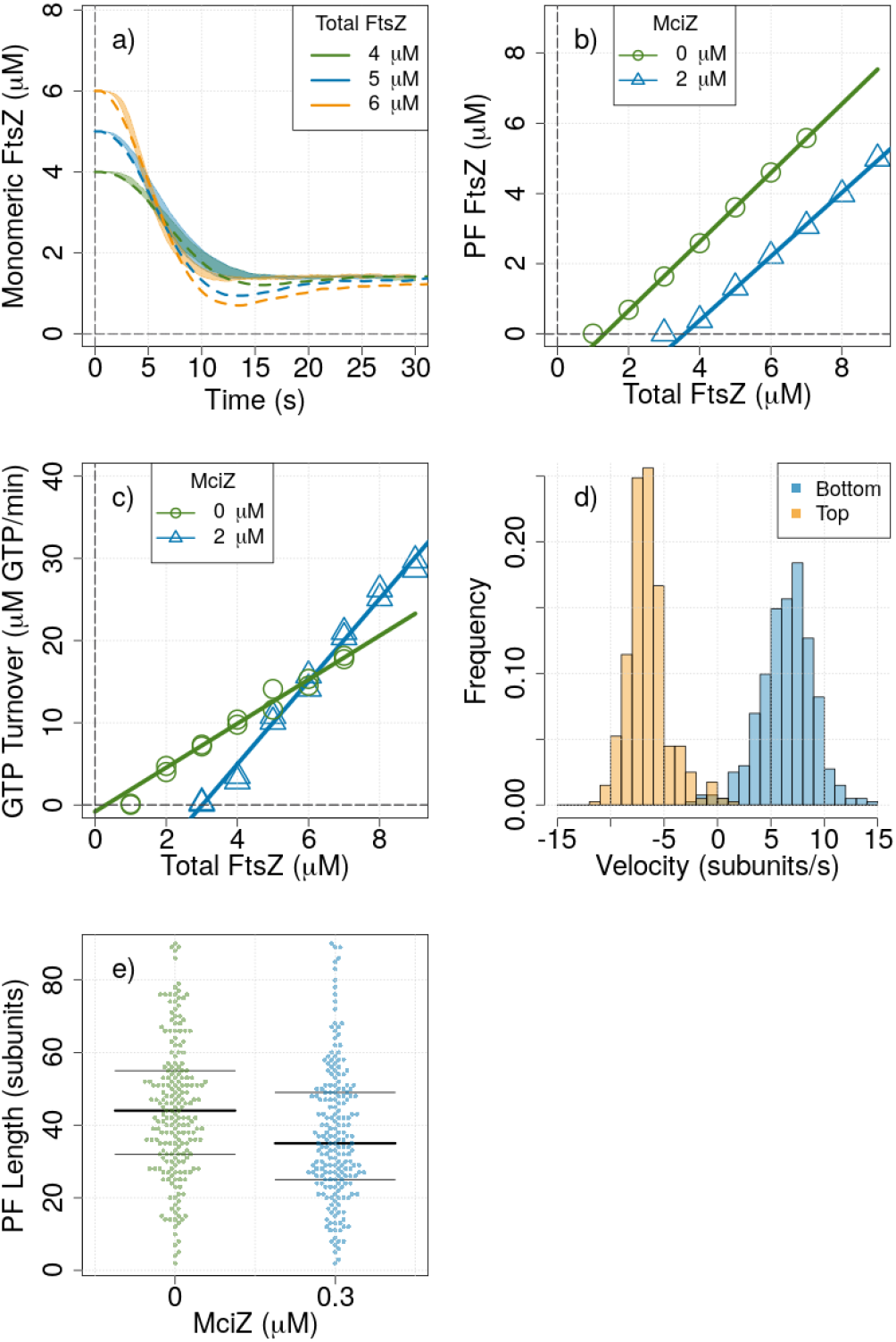
Summary of results for ***Bs*FtsZ a)** Assembly of 4, 5, or 6 μM FtsZ over 30 seconds. Dotted lines are experimental assembly data from Bisson-Filho *et al*. figure S5a. The shaded region is 95% confidence interval of 5 model runs. **b)**Concentration of subunits in the PF as a function of total FtsZ present at steady state with 0 μM MciZ (green lines and circles) or 2 μM MciZ (blue lines and triangles). The estimated C_c_ was 1.3 μM without MciZ and 3.6 μM with 2 μM MciZ. **c)** Steady state GTP turnover as a function of total FtsZ present with 0 μM MciZ (green lines and circles) or 2 μM MciZ (blue lines and triangles). Average turnover (slope of linear fit) was 2.7 GTP min^-1^ FtsZ^-1^ without MciZ and 5.0 GTP min^-1^ FtsZ^-1^ with 2 μM MciZ. Estimated C_c_ was 0.30 μM without MciZ and 3.0 μM with 2 μM MciZ. **d)** PF velocity over 5 seconds at steady state measured at PF bottom and top. The average at center is 6.7 ±2.3 subunits/s. **e)** Steady state PF length of 3 μM FtsZ with 0 μM MciZ and 0.3 μM MciZ. Avg. length was 45.3 ±20.4 subunits without MciZ and was 38.0 ±18.1 subunits with 0.3 μM MciZ.

The C_c_ predicted by the model from FtsZ in polymer was 1.3 μM (Fig. 9b, green line and circles), which fits well with experimental data (52). We also attempted to determine C_c_ from GTP turnover, as described earlier (Fig. 9e, green line and circles). Here, the C_c_ was predicted to be 0.30 μM. Again, the GTP turnover suggests a lower C_c_ than measured by assembly. The average GTP turnover (slope of line above C_c_) was 2.7 GTP min^-1^ FtsZ^-1^, which matches the reported experimental measure (52).

With these kinetics, the average treadmill velocity for all PFs was 6.7 ±2.3 subunits/s (Fig. 9d). When measured *in vivo*, mobile PFs treadmilled at approximately 6.5 ±4.0 subunits/s in *E. coli*. (11) and 7.4 ±1.8 subunits/s in *B. subtilis* (10). When we inhibited GTP hydrolysis in our model by setting k_hyd_ to zero, treadmilling ceased.

MciZ has baffled the FtsZ community since 2015 when Bisson-Filho *et. al* performed extensive experiments on *in vitro* assembly. They found that MciZ binds tightly to the bottom of the *Bs*FtsZ monomer, and the crystal structure confirmed that the bound FtsZ is in the R-state. This means that MciZ-bound subunits could function as a bottom capper by associating to the bottom of a PF and preventing further elongation. Substiochiometric concentrations (1:10) of MciZ caused a significant reduction in PF length, while higher concentrations resulted in an increase in the apparent C_c_. MciZ also increased the GTP turnover rate from 2.8 GTP min^-1^ FtsZ^-1^ to 5.4 GTP min^-1^ FtsZ^-1^ (52). We wanted to see if these observations could be explained with our model.

We modelled MciZ as a bottom capper that dimerizes with R-state monomers. This FtsZ-MciZ heterodimer then binds to the bottom of a PF with reversible kinetics and prevents further elongation. We gave the FtsZ-MciZ heterodimer the same on-off kinetics at the bottom as a FtsZ monomer.

With these parameters, we saw a decrease in PF length from 45.3 ±20.4 subunits without MciZ to 38.0±18.1 subunits with 0.3 μM MciZ with 3 μM BsFtsZ, a 1:10 stoichiometry (Fig. 9c). Experimentally, Bisson-Filho *et al*. saw a decrease in PF length from 46.5 ±17.4 subunits to 27.9 ±10.5 subunits with these same conditions with electron microscopy (52).

When we examined C_c_ based on PF FtsZ concentration vs total FtsZ concentration, we found that 2 μM MciZ increased the apparent C_c_ from 1.3 to 3.6 μM (Fig. 9f). This is somewhat higher than the experimental C_c_ of 3.1 μM, indicating that our kinetics block assembly.

We then measured GTP turnover with 2 μM MciZ. We found the GTP turnover to increase from 2.7 to 5.0 GTP min^-1^FtsZ^-1^ (Fig. 9e). This is somewhat less than the experimental measure of 5.4 GTP min^-1^ FtsZ^-1^. The increase in GTP turnover can be attributed to the increase in PF number that accompanies the decrease in PF length.

### SulA as a bottom capper or sequesterer

Finally, we applied these methods to SulA. SulA blocks PF formation both *in vivo* and *in vitro* (53–55). Crystal structures show that SulA binds to the bottom of FtsZ monomers (56). *E. coli* SulA has a moderately low affinity for FtsZ, with a K_D_ between 0.7 and 0.8 μM (55).

We are uncertain if SulA acts as a bottom capper or sequesterer. The structure of SulA suggested it could bind to the bottom of a PF and block future assembly, like a bottom capper. However, assembly and GTP turnover assays by Chen *et al*. suggested that it was a sequesterer (55). We used our model to determine if assembly assays and GTP turnover assays could differentiate between sequesterers and bottom cappers in *Ec*FtsZ.

To test this, we ran our model with SulA in equilibrium with monomeric FtsZ. To simulate a SulA capper, we allowed the FtsZ-SulA heterodimer to associate to the bottom of PFs and prevent further association. To simulate a SulA sequesterer, the FtsZ-SulA heterodimer could not associate to the bottom of PFs. In both cases, we used *Ec*F268C-FtsZ kinetics and a K_D_ of 0.78 μM for the FtsZ-SulA heterodimer.

Assembly assays based on total polymer showed an increase in C_c_ with SulA both as a bottom capper and sequesterer. The increase in C_c_ was slightly larger for the capper, but the difference was too small to confidently distinguish. Assembly assays therefore cannot distinguish between a capper and a sequesterer.

GTP turnover was more useful. Modelled as a sequesterer, SulA had no effect on the GTP turnover (Fig. 10). Modelled as a bottom capper, SulA increased the GTP turnover from 4.3 GTP min^-1^ FtsZ^-1^ to 5.3 GTP min^-1^ FtsZ^-1^. This analysis supports that SulA acts as a sequesterer.

**Figure 10:**
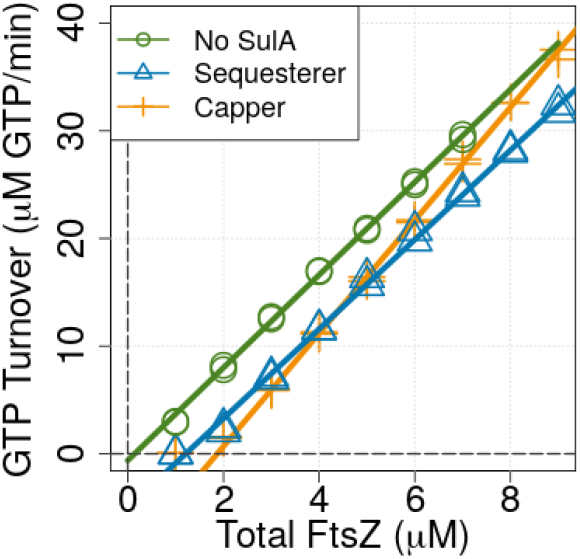
**SulA** steady state GTP turnover as a function of total FtsZ present. Model was run with either no SulA (green lines and circles), 3 μM SulA as a sequesterer (blue lines and triangles), or 3 μM SulA as a bottom capper (orange lines and crosses). The estimated GTP turnover was 4.3 GTP min^-1^ FtsZ^-1^ without SulA, 4.2 GTP min^-1^ FtsZ^-1^ with SulA as a sequesterer and 5.3 GTP min^-1^ FtsZ^-1^ with SulA as a bottom capper.

Being a sequesterer means that SulA totally inactivates the bound FtsZ. A possible mechanism is seen in an analysis of a crystal structure of *Ec*FtsZ (57). Alignment of C-terminal subdomains of R and T states showed significant movements of the T7 loop between the two forms. The structure of *Pseudamonas* FtsZ-SulA shows that SulA makes substantial contacts with FtsZ residues 206-208, in the middle of that loop (56). These contacts may lock the bound FtsZ into the R state, preventing it from switching to the T state needed for assembly.

### Potential application to actin and tubulin assembly

Actin is the classic cytoskeletal polymer that treadmills (34, 58, 59). Our model might be applied to actin, especially since recent cryoEM studies have shown a conformational change from monomer to polymer, independent of nucleotide (60, 61). However, actin assembly is complicated by the additional nucleotide state ADP-Pi (62), which has not been demonstrated for FtsZ. These states would require additional modules in the Monte-Carlo model.

We have recently suggested that the R to T transition may play a fundamental role in tubulin assembly (40). It is known from x-ray crystallography that tubulin undergoes a conformational change similar to that of FtsZ. Classical models of microtubule assembly have been been dominated by lattice models, where subunit addition at a corner is highly favored. However, a recent study concluded that all assembly steps for microtubules occur at the end of single, flared PFs, excluding corner addition (63). We have suggested that the R to T transition, which is key to FtsZ nucleation and assembly, may play a fundamental role in subunit association onto these single tubulin PFs (40). However, microtubules undergo dynamic instability, a more complex behavior than treadmilling. Integrating the R to T transition into microtubule assembly will require novel approaches.

## CONCLUSION

Overall, our model shows how the R to T conformational switch and stochastic GTP hydrolysis can drive nucleation and treadmilling and match a range of experimental observations.

## AUTHOR CONTRIBUTIONS

LC wrote the MatLab code and performed all *in silico* experiments to match experimental data. HPE suggested thermodynamic and kinetic mechanisms, and refined these in discussion with LC. LC wrote the manuscript, and HPE suggested revisions.

